# Discovery of *UPSTREAM OF FLOWERING LOCUS C* (*UFC*) and *FLOWERING LOCUS C EXPRESSOR* (*FLX*) in *Gladiolus ×hybridus*, *G. dalenii*

**DOI:** 10.1101/2021.07.02.450944

**Authors:** Jaser A. Aljaser, Neil O. Anderson, Andrzej Noyszewski

## Abstract

Gladiolus is a geophytic floricultural crop, cultivated for cut flower and garden ornamental uses. Ornamental geophytes such as gladiolus, lily, tulip and daffodil are examples of floral crops that are currently being investigated to understand the flowering pathway. While the environmental and hormonal factors leading to flowering are established in *Arabidopsis*. However, the lack of genetic regulation is poorly understood. Thus, the importance of such an ornamental crop that relies on flowers (flowering) for economic purposes encourages researchers to discover the flowering genes to breed vigorous flowering cultivars. The understanding of the flowering mechanisms in the flowering pathway is also paramount. Herein we show the discovery of *UPSTREAM OF FLOWERING LOCUS C* (*UFC*) and *FLOWERING LOCUS C EXPRESSOR* (*FLX*) genes in *Gladiolus ×hybridus and G. dalenii*. The *UFC* gene is adjacent to *FLOWERING LOCUS C* (*FLC*) which is a floral repressor in many temperate species. *FLX* gene upregulates *FRIGIDA* (*FRI*) which upregulates *FLC* expression. The discovery of both genes is a step forward in finding the *FLC* gene in gladiolus, provided they are linked. Seventeen gladiolus genotypes, consisting of early flowering and commercial cultivars, have the *UFC* gene, consisting of four exons in two allelic forms. The *UFC* gene sequenced when translated into amino acid sequence and set in pair-alignment to other species, has up to 57% in amino acid identity to *Musa acuminata*. The *UFC* protein ranges in identity with pair-alignment to other species, reaching up to 57% in amino acid identity to *Musa acuminata*. The *FLX* gene in gladiolus has 3/5 (60%) exons in relative to *Ananas comosus*, i.e. lacking 2 exons and a partially complete gene sequence; the pair-alignment of the three exons shows up over all ~65% identity of *FLX* to *Ananas comosus*. The *UFC* protein consists of a conserved domain, DUF966, which is higher in identity and pair-alignment, with up to 86% identity in *Elaeis guineensis*. The discovered *FLX* gene in gladiolus has 3/5 (60%) exons, i.e. lacking 2 exons and a partially complete gene sequence; the pair-alignment of the 3 exons shows up to ~65% of identity of *FLX* to *Ananas comosus*. These discovered two genes in gladiolus provide insight to further our understanding of the flowering and vernalization response in ornamental geophytes.

**Summary Statement:** Two gladiolus flowering genes (UFC; FLX) were discovered which will aid research in understanding flowering and vernalization in geophytes

## Introduction

“Florogenesis” is a process of flowering transitioning plant floral meristem from vegetative tissue to reproductive organs in angiosperms (Kamenetsky et al., 2012). This transition is governed by flowering genes in which expression is influenced by factors such as vernalization, photoperiod, gibberellins, autonomous pathway and ambient temperature (Srikanth and Schmid, 2011). In *Arabidopsis*, several flowering genes were discovered that are involved in flowering and act as floral integrators; some of these flowering genes are *FT*, *SOC1*, *CO*, *VRN1*, PPD, FCA, *FLD*, and *FLK* (Simpson and Dean, 2002; Simpson, 2004). These floral integrator genes specifically upregulate flowering by promoting transition from vegetative to flowering or repressing floral repressor genes. Repressor genes act as repressors of floral integrator genes and upregulates the expression of repressors, including genes such as *FLC*, *FRI*, *FLX*, *VRN2*, and SVP (Simpson and Dean, 2002; Dean et al., 2002; Ahn et al., 2007).

*FLC* and *FLC-like* is floral repressors found in many dicotyledon plants, such as *Malus* (Singh et al., 2016), *Rosa* (Zhang et al., 2017), *Coffea* (de Oliveira et al., 2014) and *Brassica* (Peacock et al., 2001). The *FLC* gene is regulated by temperature changes throughout the year, both in annuals and perennials. In summer, *FLC* expression is upregulated through *FRIGIDA* (*FRI*) by binding the *FLC* promoter through the DNA-binding protein *SUPPRESSOR OF FRIGIDA4 SUF4* (Choi et al., 2011). Also, *FRI* expression is upregulated by the *EXPRESSOR OF FLOWERING LOCUS C* (*FLX*); both *SUF4* and *FLX* are in *FRI*-specific pathway (Michaels et al., 2013). In winter, *FLC* is down-regulated through a process of vernalization as prolonged exposure of low temperature in winter in the meristem gradually reduces the expression of *FLC* (Michaels and Amasino, 1999). In addition to the vernalization pathway, the autonomous pathway reduces the expression of *FLC* both in the meristem and leaves (Michaels and Amasino, 1999). Gradual reduction of *FLC* allows *FLOWERING LOCUS T* (*FT*) to be expressed in the leaves and transported through phloem to reach to the meristematic tissue to stimulate the MADS box genes, which thereby induces flowering in *Arabidopsis* (Amasino, 2005).

In wheat and barley, the flowering pathway is regulated by photoperiod, vernalization and the circadian clock (Turner, 2013). *VRN2* acts a flowering inhibitor and only through vernalization is the expression of it downregulated. Then, the floral integrator leads to flowering in winter wheat while spring wheat doesn’t require vernalization as the *VRN2* gene is nonfunctional, although vernalization will speed up flowering in spring wheat, so vernalization acts as a facultative stimulus (Dubcovsky, 2004). In contract, maize and rice rely on plant age to build up sufficient energy requirements in order to transition to flowering through epigenetic of miR172 (Helliwell, 2011). In monocotyledon geophytes (defined as herbaceous perennial plants with underground storage organs, e.g. bulbs, corms, tubers, etc., that promote winter survival) such as *Gladiolus*, *Lilium*, *Tulipa*, *Narcissus* and *Crocus* the flowering process is poorly understood. Factors of plant growth influencing flowering in commercial geophytes are well known (Ehrich, 2013) and include photoperiod, light intensity, autonomous pathway, gibberellins, ambient temperatures and cool temperatures (vernalization; Kamenetsky, et al., 2012). On the other hand, the clear genetic pathway is still in the early stages of the discovery and characterization, in comparison to the *Arabidopsis* model which can be applied to temperate dicotyledon plants (Fadón et al., 2015). Only a few flowering genes have been discovered in geophytes (Kamenetsky, et al., 2012), such as *FT-like* in *Allium cepa* (Taylor 2009; Taylor et al 2010), *FT* in *Narcissus* (Noy-Porat, 2009), *NLF* in *Narcissus* (Noy-Porat, 2010) and *LFY* in *Allium sativum* (Rotem et al., 2007, 2011), *LFY* in *Lilium* (Wang et al., 2008). Recently, many flowering genes have been discovered in *Lilium ×formolongi*, including *FT*, *CO-like*, *AP2*, *GA1* and *SOC1* (Jia et al., 2017). The discovery of flowering genes in geophytes serve as valuable resources to draw the model pathway of flowering geophytes.

Geophytes such as *Gladiolus*, *Lilium*, *Tulipa*, *Narcissus* and *Crocus* are floricultural crops with ornamental value wherein flowering is essential to maintain the marketing value for these crops. *Gladiolus ×hybridus* Rodigas, commonly known as gladiolus, is commercially cultivated as a cut flower and as garden or landscape planta. Gladioli are geophytic plants with underground modified stems called corms, producing cormels as a means of vegetative propagation (Cohat, 1993). Flower formation is a crucial step for its success as a cut flower. Therefore, understanding the flowering pathway is vital to improve the floral market value.

Gladiolus has a genome size of 1100 Mbp, although it is unclear it is for haploid or diploid and the species is unknown (Kamo, et al., 2012), although the genome weight for gladiolus is recently measured, in *G*. *communis* 0.67-0.68 pg for monoploid G.s. (1Cx, pg) and *G*. *italicus* 0.61 pg for monoploid G.s. (1Cx, pg) (Castro, 2019). The limited knowledge in gladiolus genome is also reflected in lack of knowing gladiolus flowering genes, there is no flowering genes discovered in gladiolus except of gibberellin receptor gene *GID1a* in gladiolus (Luo et al., 2016) yet the relationship of this gene with flowering isn’t established. In *Arabidopsis*, gibberellin binds to the gibberellin receptor forming *GID1* complex that binds to *DELLA* and causing it’s degradation, enabling *SOC1* and *LFY* to upregulate, leading to flowering in the gibberellin pathway (Blázquez and Weigel, 2000; Lee et al., 2003).

Gladiolus is a monocotyledon with both summer and winter flowering species, *FLC* was not identified. It’s been hypothesized that there is no *FLC* gene in any monocotyledon species, until recently reported that *FLC* homologue were discovered in some cereal such as wheat (Sun, 2006), barley (Monteagudo, 2019) and *Brachypodium distachyon* (Kaufmann, 2013). Although the *FLC* homologue in cereals did not discover any *FRI* gene, which upregulated *FLC* expression in *Arabidopsis thaliana* (Choi et al., 2011). A hypothesis to test would be that some monocots do not possess the flowering repressor *FLC* gene and rely on alternative gene(s) to acts as a repressor(s) *miR172* through epigenetic in plant age-dependent of *Zea mays* and *Oryza* (Helliwell, 2011). Yet the question remains whether there is *FLC*-dependent pathway in all monocot plants or whether monocots are independent of *FLC*.

The *FLC* gene is located between two flanking genes, *UPSTREAM OF FLOWERING LOCUS C* (*UFC*; a gene found 4.7 Kb of upstream of *FLC*) and *DOWNSTREAM OF FLOWERING LOCUS C* (*DFC*; found 6.9 Kb of downstream of *FLC*) in *Arabidopsis* (Finnegan et al., 2004). *UFC* gene expression is repressed by vernalization, independent of *FLC* repression by vernalization (Finnegan et al., 2004). Thus, both *FLC* and *UFC* are repressed by vernalization, yet both are not dependent on each other expression; the suppression is through chromatin modification in epigenetic manner (Finnegan et al., 2004). The *VRN1* gene is expressed with vernalization and acts as a floral integrator whereas the *UFC* gene is repressed and required by *VRN1* expression dependently (Sheldon et al., 2009). The role for *UFC* in flowering is yet to be uncovered and it may not involve flowering to begin with, because vernalization only represses *UFC* in seed while *DFC* is repressed by vernalization of the plant (Sheldon et al., 2009). Insertion of the *NPTII* gene between the *UFC* and *FLC* region confirmed *NPTII* response to cold as the whole cluster region of *FLC* response to cold (Finnegan et al., 2004). In *UFC* protein, a conserved domain DUF966 is present in *Arabidopsis thaliana* which has 92 amino acid its function is still unknown (Yoshida and Weijers et al., 2019). This lack of knowledge in DUF966 function creates a challenge to identify the function of *UFC* protein. However, a recent study shows the role of *UFC* in *Arabidopsis thaliana*, the gene is designated as *SOK2*, and it would appear that it has a role in embryogenesis, root initiation, growth and branching of the primary and lateral roots (Yoshida et al., 2019). The conserved domain DUF966 is reported to be present in different species such as *Oryza sativa*, the *OsDSR* gene family contains DUF966 (Chengke and Lei, 2017). The *ZmAuxRP1* gene which promotes the biosynthesis of indole-3-acetic acid (IAA) to increase resistance against pathogens in *Zea mays* (Ye et al., 2019). The promoter for genes that contain DUF966 have a defense-stress response to pathogens, Salicylic acid, Jasmonate acid, or drought or salinity (Ye et al., 2019). Thus, all these genes that contain DUF966 vary in function but all of them are triggered by environmental or stress stimuli to overcome an undesirable change influenced plant growth and development.

The *FLX* is a gene encoding a putative leucine zipper domain that are required for *FRI*-mediated activation of *FLC* in *Arabidopsis* (Dennis et al., 2008) (Fig. 1). Expression up-regulating *FLC* occurs in winter annual *Arabidopsis* (Lee and Amasino, 2013), while late flowering phenotypes exhibit strong expression of *FLX. flx* have early flowering which indicate a role of *FLX* in suppression of flowering (Dennis et al., 2008). Several genes have been discovered in the *FLX* gene family, such as *FLX-LIKE1* (*FLL1*), *FLX-LIKE2* (*FLL2*), *FLX-LIKE3* (*FLL3*), *FLX-LIKE4* (*FLL4*) (Choi et al., 2011; Lee et al., 2013). *FLX* and *FLL4* are the most crucial genes in flowering time control in *Arabidopsis* (Lee and Amasino, 2013).

**Fig. 1.**
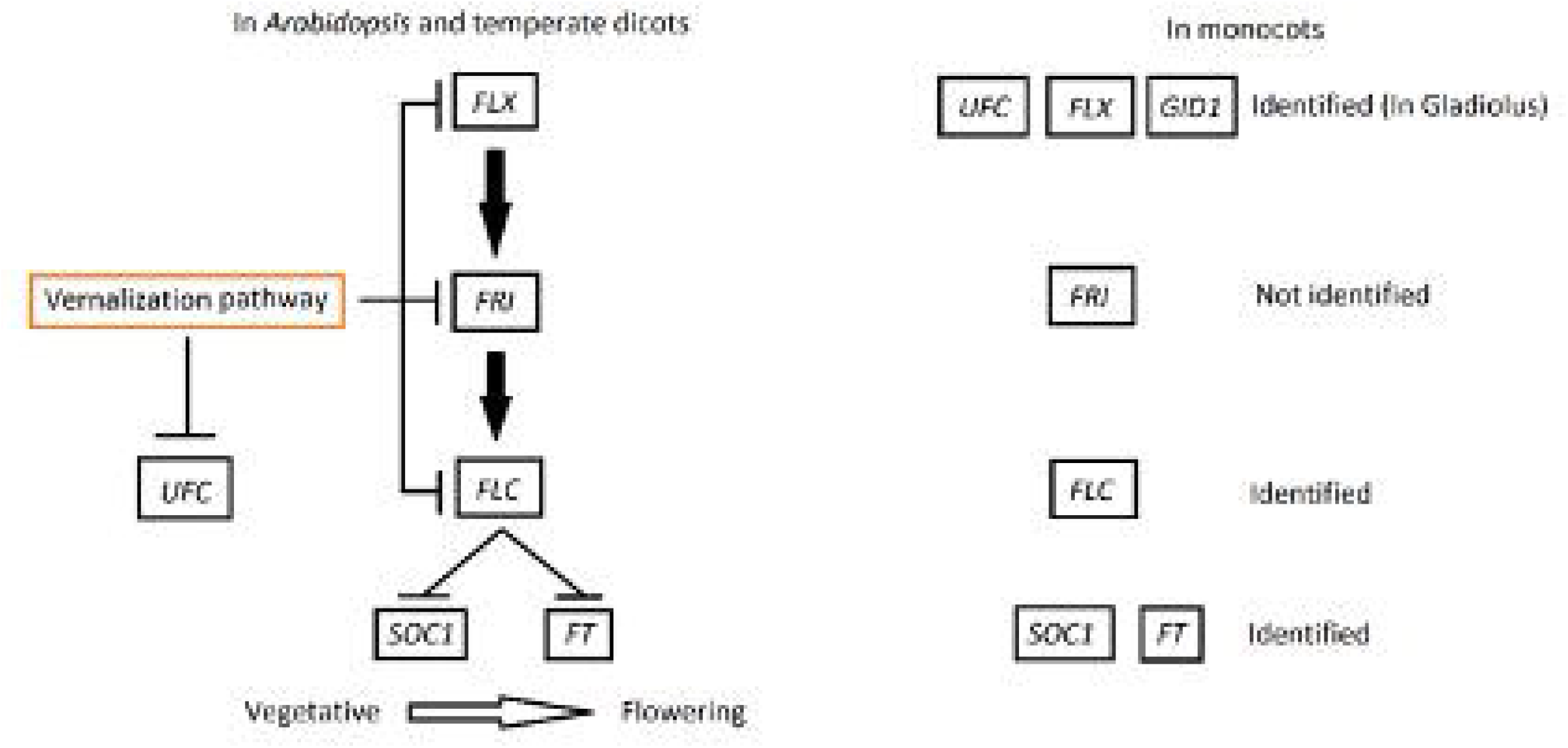
Model represents portion of flowering pathway regarding the role of *FLX* gene in flowering along with *UFC* gene in *Arabidopsis* and temperate dicots. *FLX* upregulates *FRI* which also upregulates the expression of the *FLC* protein and suppresses flowering by repressing the expression of the floral integrator *SOC1* and *FT*. Vernalization pathway downregulates the expression of *FLX*, *FRI* and *FLC* genes allowing the floral integrators to initiate flowering in vegetative state of dicots, while the vernalization pathway also downregulates the *UFC* gene (Finnegan, 2004). In monocots, *FLX*, *FLC*,*SOC1* and *FT* have been identified (Noy-Porat 2009; Amasino and Michaels 2010; Jia et al 2017; Monteagudo, 2019), while *UFC* and *FLX* have just been identified in gladiolus (in the current experiments). However, *FRI* was not identified either by lacking the presence of these repressor genes or monocots relying on other options of the flowering pathway genes.

In order to test whether *FLC* is present in gladiolus, the adjacent gene (*UFC*) will also be searched for, along with FLX which is part of the *FLC*-dependent mechanism. Therefore, the objective of this study is to identify whether *UFC* and *FLX* genes occur in gladiolus germplasm present in the University of Minnesota Gladiolus Breeding Program. The null hypotheses tested are: H_o_1 = There is no difference among gladiolus genotypes in the existence of the *UFC* gene; H_o_2 = There is no difference among gladiolus genotypes in the presence of *FLX*.

## Materials and Methods

### Germplasm Tested

The number of gladiolus genotypes used for this study is 17 (Table 1) were chosen to represent a range of diversity within cultivated gladioli (both *Gladiolus ×hybridus*, and a wild species hybrid of *G. dalenii*) which includes nine genotypes of Rapid Generation Cycling-1 (RGC-1; ones that flower in ≥1 year from seed; Anderson, 2015; Anderson, 2019; Aljaser and Anderson, 2020; Anderson and Aljaser, 2020) and eight genotypes of Non-RGC (that require >1 to 5 years to flower from seed). Fourteen of these genotypes are parents and hybrids created by the University of Minnesota Gladiolus Breeding Program, while three additional genotypes are commercial cultivars. One genotype ‘Carolina Primrose’ is derived from the species *G. dalenii* (Table 1). All gladiolus pedigrees used in this experiment are published (Aljaser and Anderson, 2020; Anderson and Aljaser, 2020) and commercial cultivars ‘Beatrice’ (an open-pollinated seedlings of unknown origin, occurring in a private garden, Brookfield, Vermont, in 2003. This genotype was selected for its winter hardiness, surviving in USDA Z3). ‘Glamini’^®^ a series of shorter in height than tall summer gladiolus, bred by Dutch breeders, bloom early, has a range of flowering colors (Wayside Gardens, 2020). ‘Carolina Primrose’ is an heirloom gladiolus, bred in 1908, yellow color flowers, collected at an old homesite in North Carolina and it is a cultivar bred from *G. primulinus* (Old House Gardens, 2020).

**Table 1.** The *Gladiolus* genotypes used in this study, their codes and whether they are Rapid Generation Cycling (RGC): + is for RGC genotypes and for Non-RGC genotypes; all gladiolus genotypes were tested for the presence of *UFC* gene, *FLX* gene and *FRI* gene.

| Genotype | Code | RGC |
| --- | --- | --- |
| *Gladiolus* ×*hybridus* 21213 | 1 | + |
| *Gladiolus* ×*hybridus* 2220 | 2 | + |
| *Gladiolus* ×*hybridus* 2231 | 3 | + |
| *Gladiolus* ×*hybridus* 2337 | 4 | + |
| *Gladiolus* ×*hybridus* 35314 | 5 | + |
| *Gladiolus* ×*hybridus* 3923 | 6 | + |
| *Gladiolus* ×*hybridus* 3931 | 7 | + |
| *Gladiolus* ×*hybridus* 74210 | 8 | + |
| *Gladiolus* ×*hybridus* 7736 | 9 | + |
| *Gladiolus* ×*hybridus* 28236 | 10 | - |
| *Gladiolus* ×*hybridus* 15531 | 11 | - |
| *Gladiolus* ×*hybridus* 20732 | 12 | - |
| *Gladiolus* ×*hybridus* 60314 | 13 | - |
| *Gladiolus dalenii* ‘Carolina Primrose’ | 14 | - |
| *Gladiolus* ×*hybridus* ‘Beatrice’ | 15 | - |
| *Gladiolus* ×*hybridus* ‘Glamini’® | 16 | - |
| *Gladiolus* ×*hybridus* ‘Vista | 17 | - |

### Greenhouse Environment

Mature (capable of flowering) gladiolus corms were planted into 1679.776 cm^2^ square, deep pots (Belden Plastics, St. Paul. MN) in week 23 (2017) and grown for 18 weeks. Containers were filled with SS#8-F2-RSi potting soil, “SunGrow” (Sun Gro Horticulture, Agawam. MA). The corms were grown in a long day photoperiod (0800 – 1600 HR supplied by 400-W high-pressure sodium lamps + 2200 to 0200 HR night interruption, >150 μmol m^-2^ sec^-1^) at a minimum setpoint of 18° C (day/night), 70-80% relative humidity, with irrigation accomplished using constant liquid feed (CLF) of 125 ppm N from water-soluble 20N–4.4P–16.6K (Scotts, Marysville, OH) and deionized water on weekends. Standard fungicide drenches and insecticides were applied either monthly or as needed, respectively.

### DNA extraction and probe design

Newly expanded gladiolus leaves were harvested, placed in an ice box and sent to RAPiD Genomics^®^ LLC (Gainesville, FL; http://rapid-genomics.com/home/) for DNA extraction, probe design, sequencing and computable analysis. Probe designs for the *UFC* gene were based on banana, *Musa acuminata* subsp. *malaccensis* accession XM_009383889, from the GenBank Nucleotide Core (NCBI, 2016a) and oil palm, *Elaeis guineensis* accession XM_010920607.2 (NCBI, 2017a). Probe design for *FLX* gene were based on oil palm, *Elaeis guineensis* accession XM_010924316.2 (NCBI, 2017b) and date palm, *Phoenix dactylifera* accession XM_008801571.2 (NCBI, 2016b). The designed probe for *UFC* able to capture the locus in *Musa acuminata* and *Elaeis guineensis* by capturing the 2x coverage of the *UFC* exons in *Musa acuminata* and *Elaeis guineensis*. While the *FLX* probe capture the locus in *Elaeis guineensis* and *Phoenix dactylifera*. Then the probes able to amplify short read of *UFC* and *FLX* genes in gladiolus. The reads are sequenced through Illumina dye sequencing technique, the raw data is demultiplexed using Illuminas BCLtofastq then assembled using MaSuRCA^®^ software (Zimin et al., 2013) creating full assembly sequences scaffolds. Afterword, read mapping using reference genome and blast to filter all assembled sequences for hits to the sequences provided for probes design (*UFC* and *FLX*), then count read numbers for each assembled sequence that passed the filter and accruing the final sequences for genetic analysis. Gene sequences will be deposited into GenBank.

### Genetic Analysis

The sequence data for the *UFC* and *FLX* genes used in this study were found in the genetic sequence database under the following accession/ID numbers: *Ananas comosus* (Aco009327) *UFC* gene is from the Pineapple Genomics Database (Yu, et al., 2018); *Musa acuminata* (GSMUA_Achr5T28540_001) *UFC* from the Banana Genome Hub (Droc et al., 2013); *Elaeis guineensis* (p5.00_sc00099_p0095) *UFC* from the Malaysian Oil Palm Genome Programme (Halim et al., 2018); *Asparagus officinalis* (evm.model.AsparagusV1_08.3493) *UFC* from the Asparagus Genome Project (Harkess et al., 2017); *Arabidopsis thaliana* (At5g10150) *UFC* from The Arabidopsis Information Resource (TAIR) (TAIR, 2020); *Glycine max* (Glyma.11G193000.1) *UFC* from the SoyBase (Grant et al., 2010). The *FLX* protein was from the GenBank Nucleotide Core with accession numbers as follows: *Ananas comosus* (XP_020095672.1) (NCBI, 2017c), *Musa acuminata* (XP_009420070.1) (NCBI 2016c), *Elaeis guineensis* (XP_010922618.1) (NCBI, 2019a), *Arabidopsis thaliana* – *FLX* (NP_001154541.1) (NCBI 2019b), *Arabidopsis thaliana* – *FLL1* (NP_566492.1) (NCBI 2019c), *Arabidopsis thaliana* – *FLL2* (NP_001320766.1) (NCBI 2019d), *Arabidopsis thaliana* – *FLL3* (NP_564678.1) (NCBI 2019e), *Arabidopsis thaliana* – *FLL4* (NP_001119474.1) (NCBI 2019f) and *Glycine max* (Glyma.15g269300) *FLX* was from the SoyBase (Grant et al., 2010).

Generated sequences were analyzed for gene prediction using the HMM-based gene structure prediction of FGENESH with *Arabidopsis thaliana* (Generic) as the specific gene-finding parameter. The predicted genes for *UFC* and *FLX* were analyzed in multi-alignment using Geneious^©^ software (Biomatters Ltd, Auckland, NZ). The *UFC* protein sequences of gladiolus were analyzed for conserved domains using the Protein Homology/analogy Recognition Engine V 2.0 (Phyre2) browser (Kelley et al., 2015). Then the alignment of conserved domain was formed to compare the matching and differences in each amino acid in the sequences of gladiolus and the other comparison species. A phylogenetic tree of all *UFC* genotypes of *Gladiolus* was formed by computing the distances using the Tamura-Nei method and were in the units of the number of base substitutions per site. The tree building used the Neighbor-Joining method and a bootstrap test was performed for each tree (500 replicates).

## Results

The investigated genotypes of *Gladiolus* resulted from the gladiolus breeding program for the selection of rapid generation cycling, which includes 14 breeding genotypes and 3 commercial cultivars (‘Vista’, ‘Glamini^®^’ and ‘Carolina Primrose’; Table 1). Rapid Generation Cycling (RGC) in gladiolus is the ability of flowering in the first year or less form seed sowing (Aljaser and Anderson, 2020; Anderson and Aljaser, 2020). Typically, gladiolus are juvenile in the first few years and require 3-5+ years to reach reproductive age. The University of Minnesota Gladiolus breeding program developed certain gladiolus genotypes able to flower in the first year of seed sowing (Aljaser and Anderson, 2020; Anderson and Aljaser, 2020).

The designed probe for the *UFC* gene in *Gladiolus* is done by RAPiD genomics^®^ LLC (Gainesville, FL; http://rapid-genomics.com/home/) resulted in the generation in total of 433 sequences; 161 sequences among them had read hits of the *UFC* gene with various percentage of coverage. Of these, 34 selected sequences were chosen for this study, based on the largest length with two sequences per genotype due to presence of two alleles per genotype. These sequences represent the genomic sequence of *UFC* in gladiolus. The sequences were analyzed for gene prediction using the HMM-based gene structure prediction of FGENESH by having *Arabidopsis thaliana* (Generic) as the specific gene-finding parameter as the gene prediction is optimized for *Arabidopsis thaliana*, the results confirmed the presence of *UFC* exons of a protein and coding sequence. The *UFC* coding sequences of each gladiolus genotype were analyzed in Genoeious^®^ by pair-alignment with its genomic sequence to determine each exon. After the pair-alignment, the Coding sequence was then translated and aligned in multi-alignment process using MUSCLE alignment in the neighbor joining clustering method and CLUSTALW sequencing scheme with *UFC* proteins of other species: *Ananas comosus*, *Musa acuminata*, *Elaeis guineensis* and *Asparagus officinalis*, *Arabidopsis thaliana* and *Glycine max*. The *UFC* gene in *Gladiolus ×hybridus* was assigned to *GhUFC* as a label while genotype 14, *G. dalenii* ‘Carolina Primrose’, is assigned to *GdUFC*. In terms of the *FLX* gene, *Gladiolus ×hybridus* is assigned to *GhFLX* and genotype 14 *G. dalenii* ‘Carolina Primrose’ is *GdFLX*. There are two alleles of the *UFC* gene found in gladiolus. The designated alleles are A and B. Thus, the genes are designated as *GhUFC-A* and *GhUFC-B*. The median number of amino acids of *GhUFC-A* protein is 420 amino acids across all gladiolus genotypes, with some genotypes having less than 420 amino acids and one genotype has 451 amino acids which could be due to an insertion, while *GdUFC-A* also has 420 amino acids. The second allele, *GhUFC-B* protein has a range of 375 to 410 amino acids, *GdUFC-B* has 286 amino acids with an incomplete protein, missing many amino acids and a stop codon (Figs 2, 3). Gladiolus genotypes 1 and 15 were selected for pair-alignment with the species of comparison (Table 2), the table shows similarities of identity protein sequences and the percentage of *GhUFC-A* and *GhUFC-B* occur at range of ~30% to 57% across all species (Table 2). The intron-exon organization of *GhUFC-A* 15 genotype is similar to *Elaeis guineensis* and *Asparagus officinalis* in term of exons splicing (Fig. 4) while *GhUFC-B* genotype 1 has some similarity with *Ananas comosus* exons splicing. The remaining genotypes fall into these two configurations of the exon; the configuration shows the location of the conserved domain for *UFC* gene DUF966, which is found in *Arabidopsis* and other selected species of comparison. The DUF966 domain has the 92 amino acid and 93 amino acid of *Ananas comosus*. The multi alignment of the *UFC* protein conserved domain is overall conserved across species, although it is polymorphic (Fig. 5). With the high identity matching in *Gladiolus* genotypes 1 and 15 exhibit a range of ~65% to ~86% across all investigated species for the DUF966 domain of the *UFC* protein (Table 3). The *GhUFC-A* allele has a high identity across gladiolus genotypes, with the polymorphic exception of genotypes 3 and 7, due to (Fig. 6). *GhUFC-B* is also conserved and identical in sequence with genotype 12, due to missing amino acids.

**Fig. 2.**
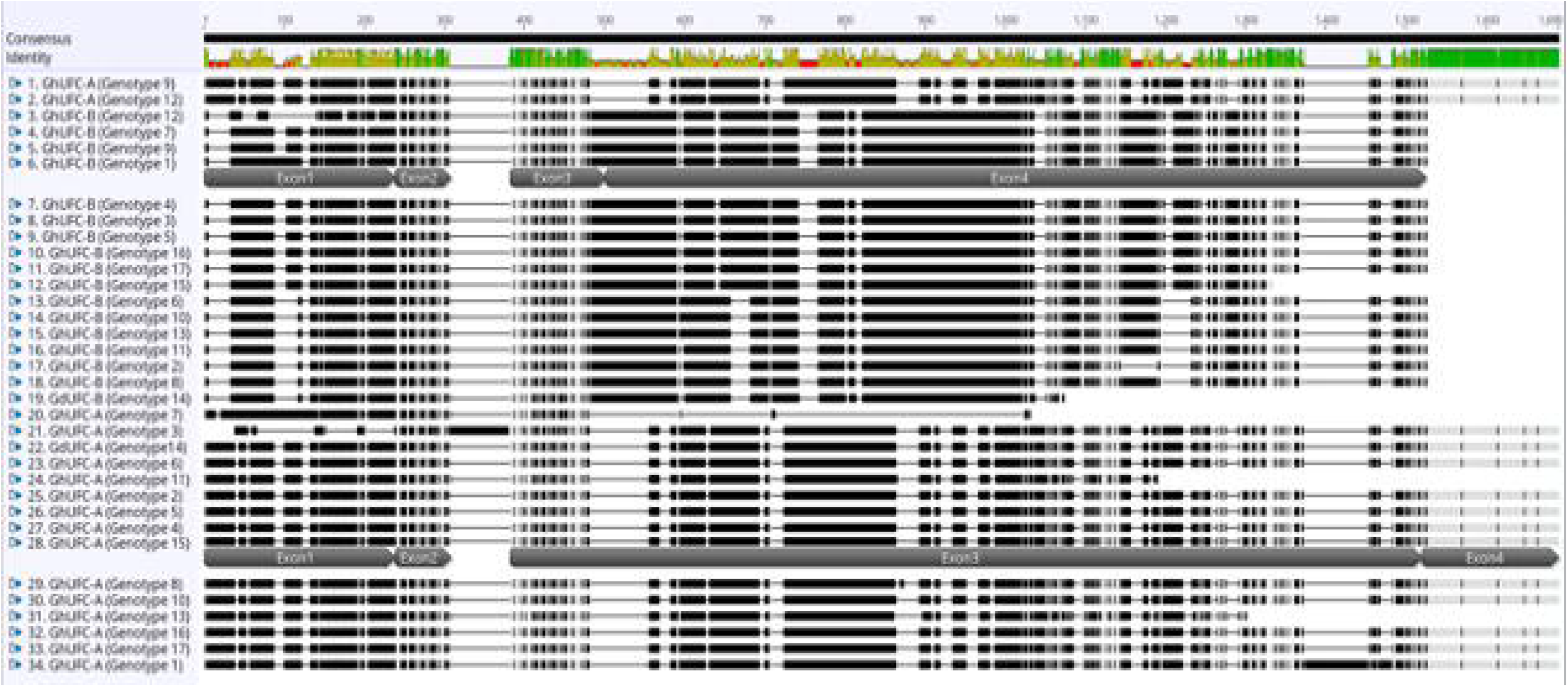
Multi-alignment of *UFC* coding sequence in *Gladiolus ×hybridus* (*GhUFC*) and *Gladiolus dalenii* (*GdUFC*), the alignment is for the 17 genotypes, each genotype has 2 alleles, allele A and allele B: *GhUFC-A*, *GdUFC-A*, *GhUFC-B*, *GdUFC-B*. Both alleles has 4 exons but allele A size is larger in coding sequence than allele B. The alignemtn shows insertion and missing coding sequences in some genotypes. The multi-alignment is done in MUSCLE pair-alignment using neighbor joining cluster method and CLUSTALW sequencing scheme (Geneious)^®^

**Fig. 3.**
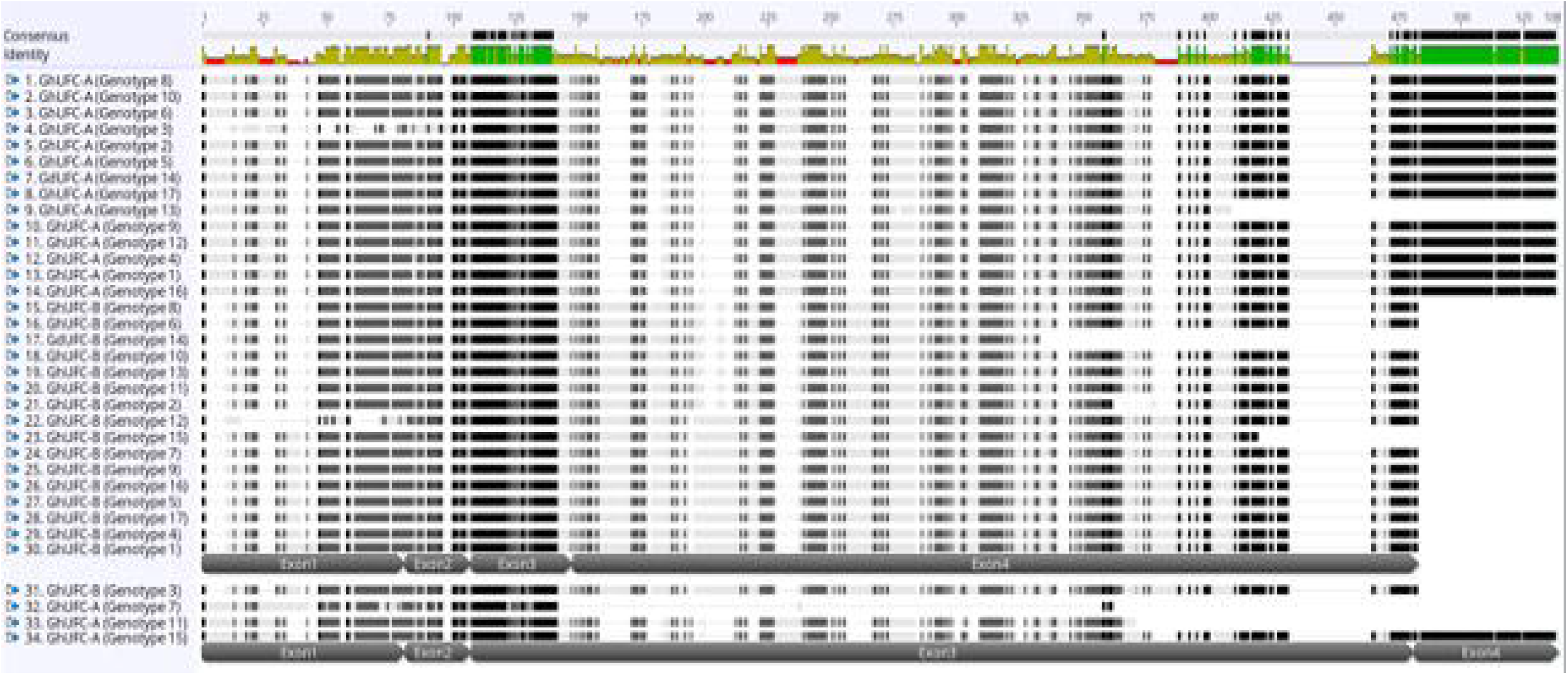
Multi-alignment of *UFC* amino acid sequence in *Gladiolus ×hybridus* (*GhUFC*) and *Gladiolus dalenii* (*GdUFC*), the alignment is for the 17 genotypes, each genotype has 2 alleles, allele A and allele B: *GhUFC-A*, *GdUFC-A*, *GhUFC-B*, *GdUFC-B*. Both alleles has 4 exons but allele A size is larger in amino acid sequence than allele B. The alignemtn shows insertion and missing amino acid sequences in some genotypes. The alighnment identify conserved amino acid sequences (green color). The multialignment is done in MUSCLE pair-alignment using neighbor joining cluster method and CLUSTALW sequencing scheme (Geneious)^®^

**Fig. 4.**
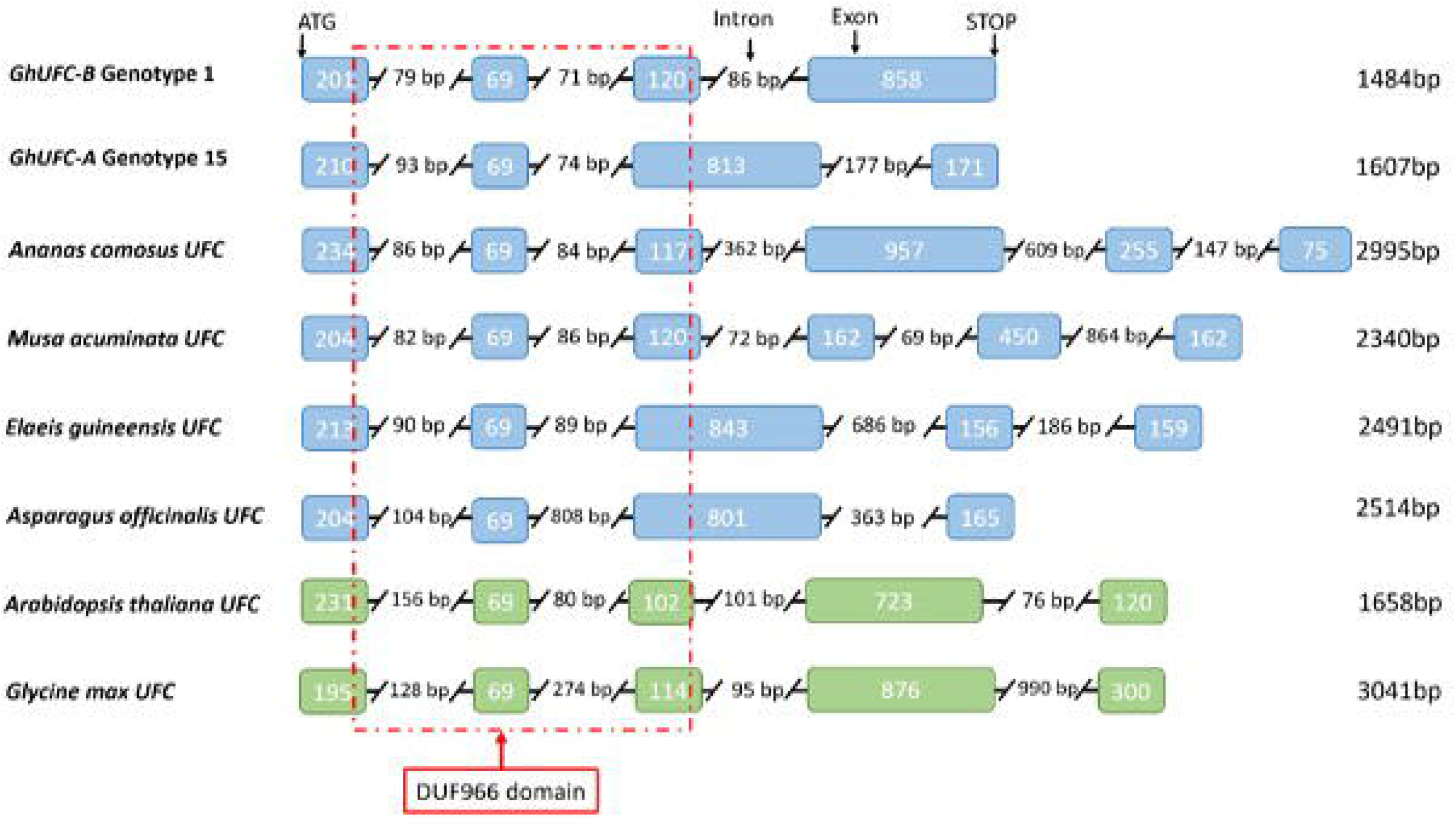
Intron-exon configuration of the *UFC* genes in *Gladiolus ×hybridus* of genotypes 1 and 15 in relation to several species. Monocot species are highlighted in blue: *Ananas comosus*, *Musa acuminata*, *Elaeis guineensis* and *Asparagus officinalis*. Dicot species highlighted in green for *Arabidopsis thaliana* and *Glycine max*. The intron-exon organization of *GhUFC-A* genotype 15 is similar to *Elaeis guineensis* and *Asparagus officinalis* in terms of exons splicing while *GhUFC-B* genotype 1 has some similarity with *Ananas comosus* exons. Sequences were aligned based on first exon sequences. Total length of the gene’s coding region is listed on the right of each respected species. The red line represents the conserved domain DUF966 in relation to the location of the domain with exon configuration.

**Fig. 5.**
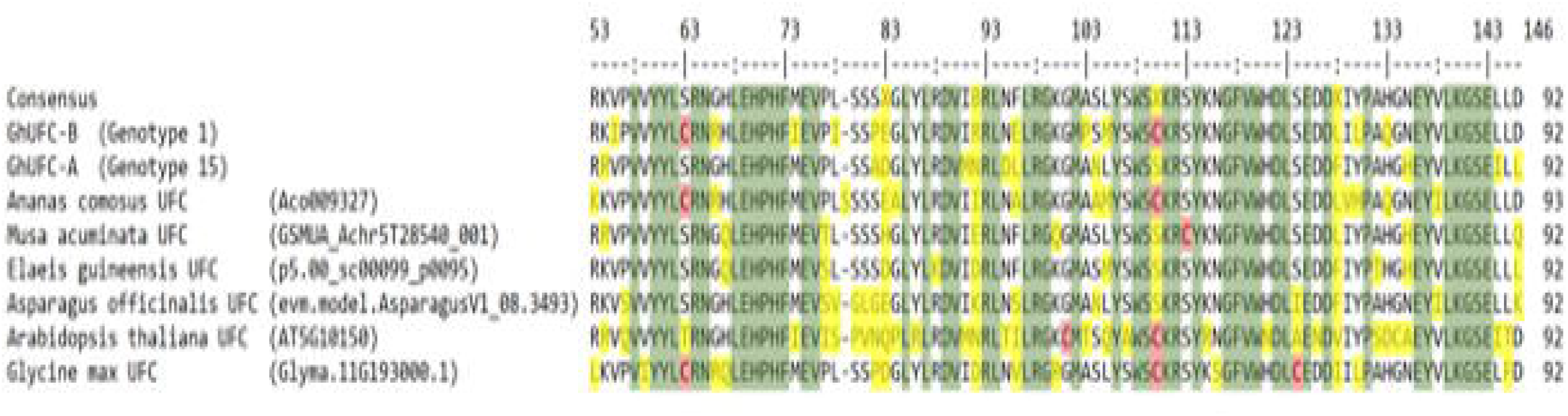
Alignment of the globular region containing DUF966 domain of UFC proteins from *Gladiolus ×hybridus* of genotypes 1 and 15, *Ananas comosus*, *Musa acuminata*, *Elaeis guineensis Asparagus officinalis*, *Arabidopsis thaliana* and *Glycine max*. Green coloration shows identical amino acid sequence; yellow color highlights the polymorphisms while red color shows the cytosine amino acid. The conserved domain DUF966 is 92 amino acids.

**Fig. 6.**
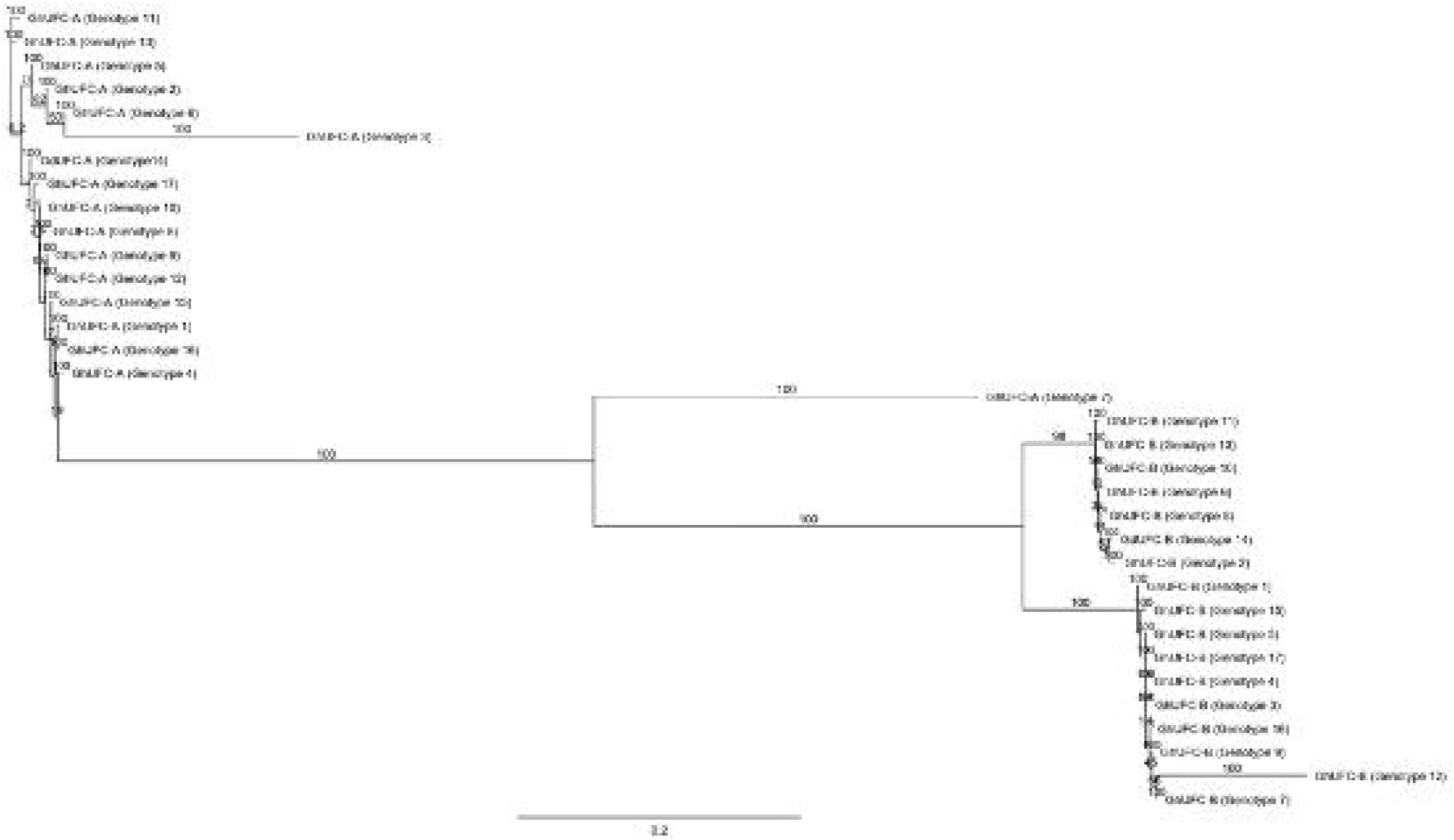
The phylogenetic tree of all *UFC* genotypes in *Gladiolus*; genetic distances were computed using the Tamura-Nei method and are in the units of the number of base substitutions per site. The tree build using the Neighbor-Joining method and the bootstrap test was performed for each tree (500 replicates) and the tree format is organized and ordered with a scale bar of 0.2 (Geneious^®^).

**Table 2.**
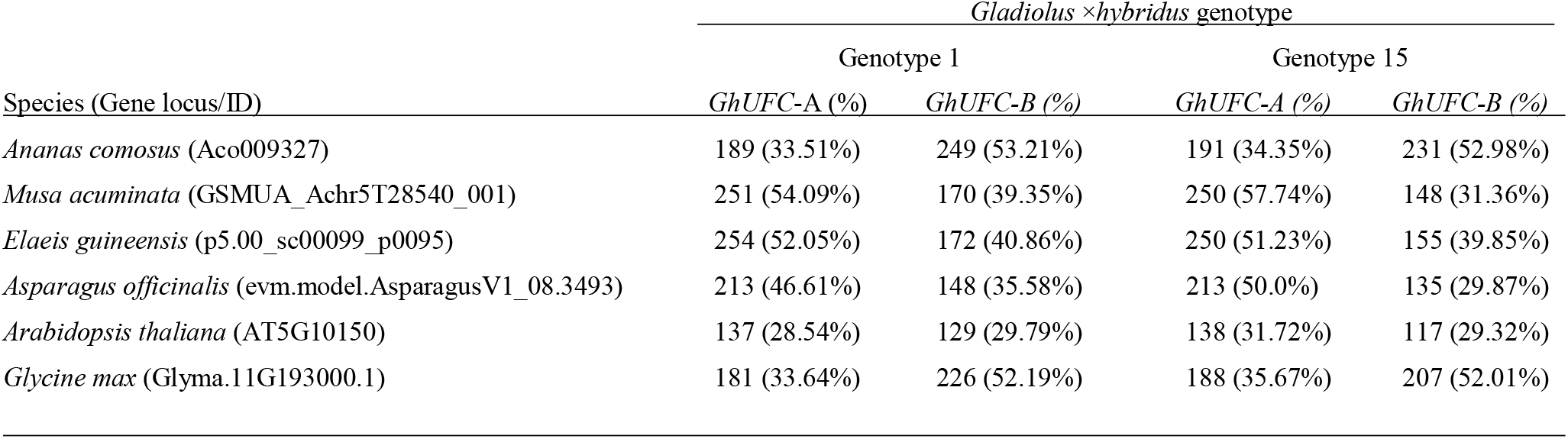
The identity of amino acid sequences and number (%) of two *UFC* proteins (*GhUFC-A*, *GhUFC-B*) in two *Gladiolus* (genotypes 1 and 15) in relation to other species (Gene locus/ID) through pair alignment; the similarities of sequences in identity of gladiolus genotypes ranges from ~30% to 57% across all investigated species for the whole *UFC* protein; alignment is done with MUSCLE alignment using the neighbor joining clustering method and the CLUSTALW sequencing scheme (Geneious^®^).

**Table 3.** Identity of amino acid sequences, number (%) of *UFC* proteins in the conserved domain DUF966 in *Gladiolus* genotypes 1 and 15 in relation to other species through pair alignment. *Gladiolus* genotypes 1 and 15 exhibit a range of ~65% to ~86% across all investigated species for the DUF966 domain of *UFC* protein.

| *Gladiolus* × *hybridus* genotype |  |  |  |  |
| --- | --- | --- | --- | --- |
| Species (Gene locus/ID) | <u>Genotype 1</u> |  | <u>Genotype 15</u> |  |
|  | *GhUFC-A* (%) | *GhUFC-B* (%) | *GhUFC-A* (%) | *GhUFC-B* (%) |
| 646 *Ananas comosus* (Aco009327) | 70 (76.09%) | 79 (84.95%) | 79 (84.95%) | 79 (84.95%) |
| 647 *Musa acuminata* (GSMUA_Achr5T28540_001) | 78 (84.78%) | 71 (78.02%) | 78 (84.78%) | 71 (78.02%) |
| 648 *Elaeis guineensis* (p5.00_sc00099_p0095) | 79 (85.87%) | 72 (79.12%) | 79 (85.87%) | 72 (79.12%) |
| 649 *Asparagus officinalis* (evm.model.AsparagusV1_08.3493) | 75 (82.42%) | 70 (76.09%) | 75 (82.42%) | 70 (76.09%) |
| 650 *Arabidopsis thaliana* (AT5G10150) | 61 (66.30%) | 61 (64.89%) | 62 (67.39%) | 61 (64.89%) |
| 651 *Glycine max* (Glyma.11G193000.1) | 70 (76.92%) | 75 (82.42%) | 70 (76.92%) | 75 (81.52%) |

*FLX* gene was identified in gladiolus, the total genotypes which had presences of *FLX* are 12 out of 17 genotypes; 11 genotypes have *GhFLX* whereas *GdFLX* is identified in genotype 14 (Table 4). The range of amino acid proteins are from 146 to 254 amino acids missing the stop codon. Three genotype sequences (genotype 3, 6 and 16) have the longest amino acid chain. *FLX* is present in many species, in *Arabidopsis* alone it belongs to *FLX* gene family, *FLX*, *FLOWERING LOCUS C EXPRESSOR-LIKE 1* (*FLL1*), (*FLL2*), (*FLL3*) and (*FLL4*) (Lee and Amasino, 2013). Based on the pair-alignment, *GhFLX* and *GdFLX* matches *FLL1* as high as 50% in amino acid identity (Table 4), the similarities of sequences in identity of gladiolus genotypes ranging from ~26% to ~65% across all investigated species of the whole *FLX* protein, the highest identity is in *Ananas comosus* match with ~65%. The multi-alignment for all *FLX* indicates that the tested gladiolus genotypes which has the longest amino acid sequences lacks exons (Fig. 7), a pair-alignment test with *Ananas comosus* – *FLX* reveals *GhFLX* genotype 16 possibly lacks two exons and stop codon (Fig. 8).

**Fig. 7.**
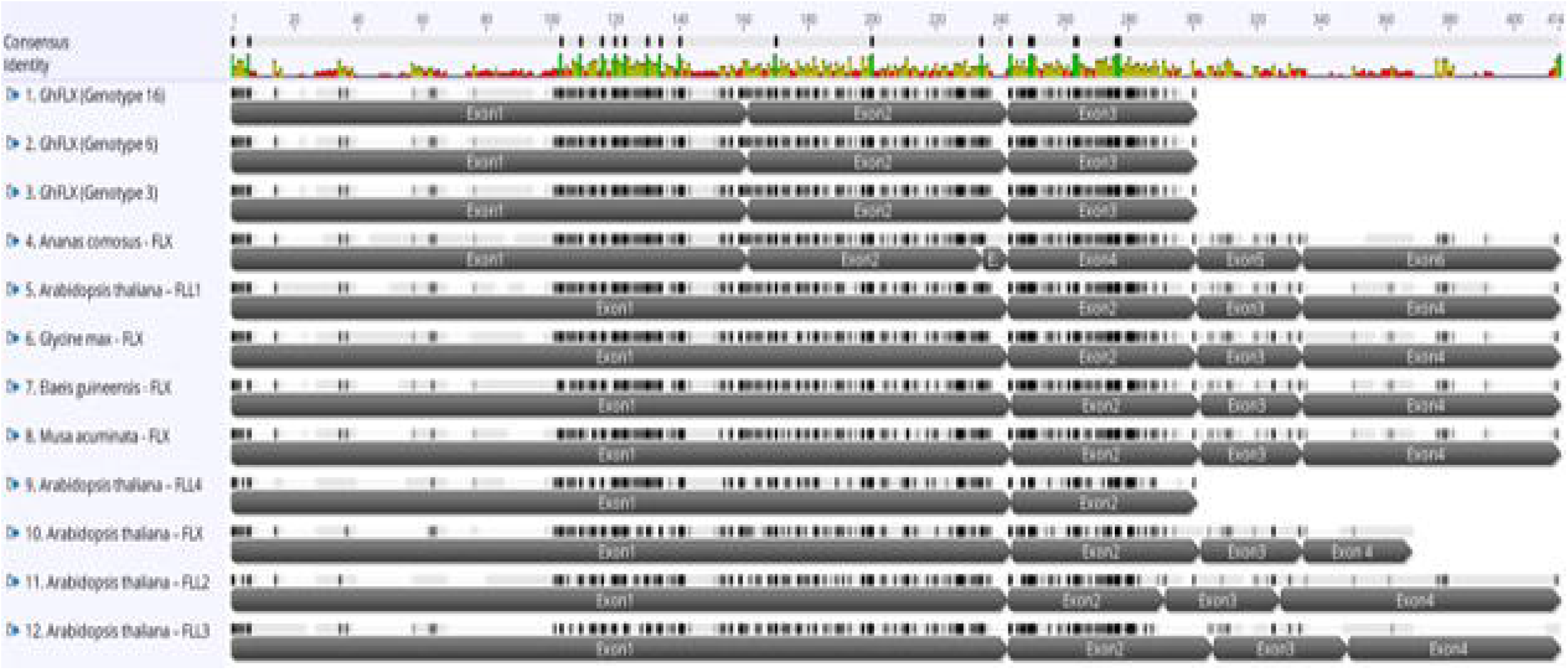
Multi-alignment of *FLX* amino acid sequence in *Gladiolus ×hybridus* (*GhFLX*) with other species; *Arabidopsis thaliana*, *Ananas comosus*, *Elaeis guineensis*, *Musa acuminata* and *Glycine max*. The alignment is for the 3 gladiolus genotypes (3, 6 and 16) each genotype has 3 exons. The alignment shows missing amino acid sequences in gladiolus genotypes 3, 6 and 16 as amino acid sequences does not have a stop codon. The alignment identifies conserved amino acid sequences (green color). Note *Arabidopsis thaliana* – *FLL4* is a functional protein which has two exons only (Lee and Amasino, 2013). The multialignment is done in MUSCLE pair-alignment using neighbor joining cluster method and CLUSTALW sequencing scheme (Geneious)^®^.

**Fig. 8.**
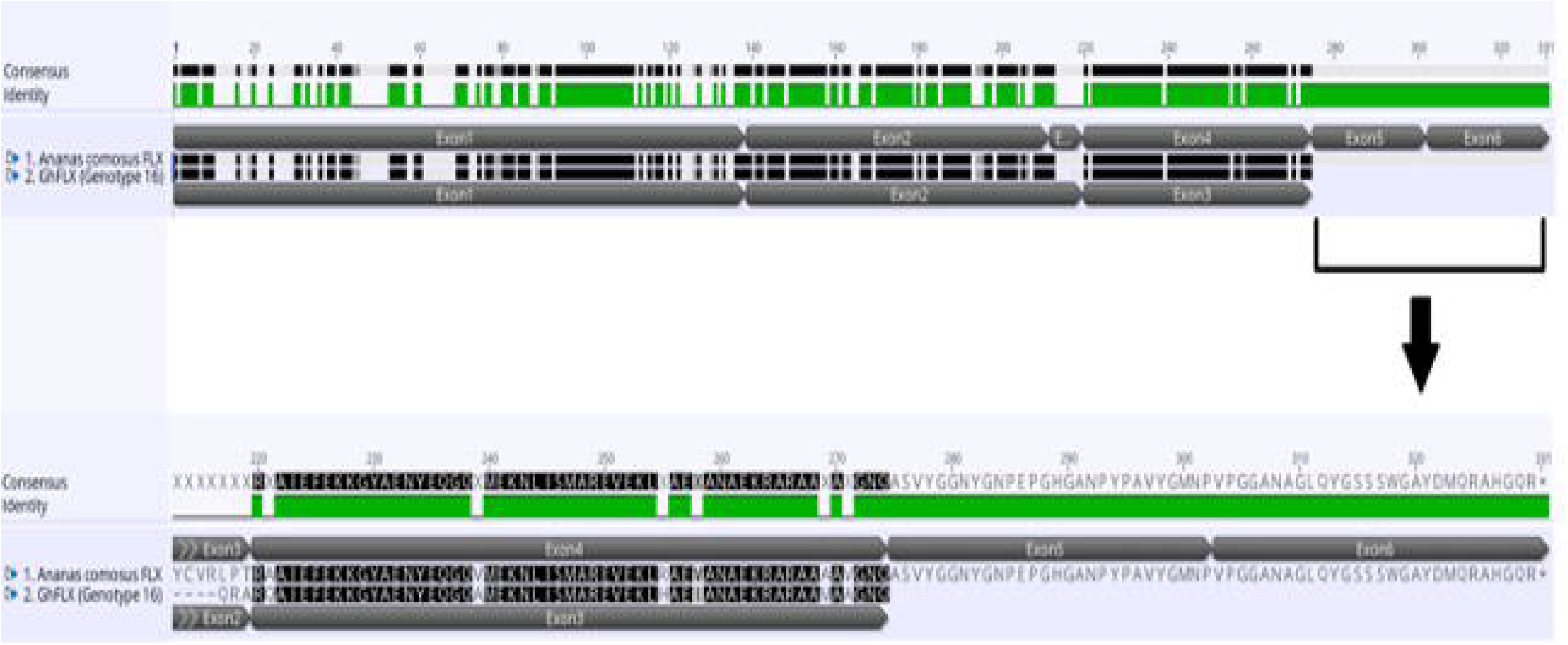
Pair alignment of *FLX* protein in *Ananas comosus* and *Gladiolus ×hybridus* genotype 16 showing the complete *FLX* protein in *Ananas comosus* with five exons while incomplete *FLX* protein in *Gladiolus ×hybridus* (*GhFLX*) which lacks the remaining two exons and stop codon. The alignment is done in MUSCLE pair-alignment using neighbor joining cluster method and CLUSTALW sequencing scheme (Geneious)^®^.

**Table 4.**
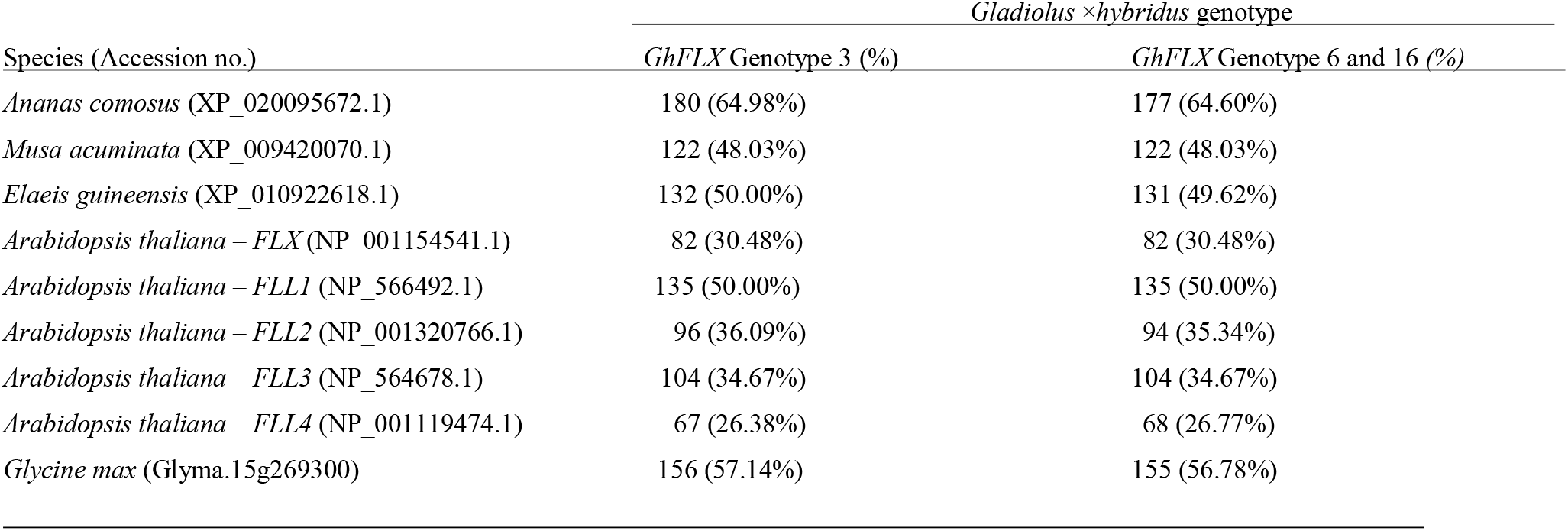
Number (%) of amino acid sequences of *GhFLX* protein in *Gladiolus* genotypes 3 and 6 (genotype 16 is identical to genotype 6) in relation to the other species through pair alignment; similarities of sequences in identity of gladiolus genotypes ranged from ~26% to ~65% across all investigated species of the whole *FLX* protein; alignment is done in MUSCLE, using the neighbor joining clustering method and the CLUSTALW sequencing scheme (Geneious^®^).

## Discussion

The presence of a putative *UFC* gene in gladiolus is confirmed with two alleles, *GhUFC-A* and *GhUFC-B*. It is highly possible that as the current gladiolus hybrids on the commercial market are mostly polyploids, primarily tetraploids with 60 chromosomes (2*n* = 4 × = 60) (Bamford, 1935; Saito and Kusakari, 1972; Ohri, 1985). Many of the gladiolus cultivars are interspecific hybrids (Benschop, 2010). Therefore, the presence of different alleles of genes is expected in the genotypes chosen for this study, as there are 13 genotypes are from University of Minnesota gladiolus breeding program and the other 4 genotypes are from commercial gladiolus cultivars (Table 1). *GhUFC-A* gene has relatively ~50% identity with *Musa acuminata*, *Elaeis guineensis* and *Asparagus officinalis* (Table 2), the splicing of *Elaeis guineensis* and *Asparagus officinalis* exons is similar to *GhUFC-A* (Fig. 4). *GhUFC-B* gene splicing is similar to *Ananas comosus*, *Arabidopsis thaliana* and *Glycine max UFC* gene splicing in the first 4 exons (Fig. 4). These divergence in splicing of the *UFC* gene support the *UFC* gene being present in gladiolus with two alleles due to the possible of ploidy level, as many of gladiolus commercial are polyploids (Bamford, 1935). Further tests should be done to identify whether or not the *UFC* gene is also present in diploid gladiolus species, such as *G. murielae*, *G. tristis* and *G*. *carneus* since these three species only exist in the diploid form (Goldblatt and Manning, 1998).

The *UFC* gene is responsive to vernalization by lowering expression alongside *FLC* and *DFC* in *Arabidopsis thaliana*, as all these genes are in the cluster of vernalization stimulus region (Finnegan et al., 2004). *FLC* is a floral repressor, the overexpression of *FLC* results in a delay in flowering (Amasino, 1999), while overexpression of *UFC* does not result in the altering flowering time (Finnegan et al., 2004). Thus, *UFC* is adjacent to *FLC* both repressed by vernalization, yet *UFC* does not show any influence in flowering time. This was observed herein because the genotypes for this study are a mixture of RGC-1, which are early flowering gladiolus able to reach flowering in the first year from seed and the classical later-flowering gladiolus which requires 3-5+ years to flower from seed. The multi-alignment of *UFC* protein of RGC-1 does not show any difference from non-RGC-1 genotypes, thus the *UFC* gene most likely isn’t involved in flowering, directly at least and this was proven in a *UFC* study in *Arabidopsis thaliana* (Sheldon et al., 2009). The main differences are shown in Fig. 6 which represents the differences between alleles of *UFC-A* and *UFC-B*, regardless of the gladiolus genotypes tested herein (Fig. 6).

In winter, vernalization suppress both *FLC* and *UFC* expression (Finnegan 2004), which allows floral genes integrators to promote flowering with the hypothesis would be that after vernalization and flowering, the *UFC* protein involvement is in embryogenesis and root initiation such that growth and branching occur in the spring season (since it doesn’t occur in the winter season). This could explain how both *FLC* and *UFC* are both negatively responsive to vernalization stimuli in the cluster genes area, while upstream of *UFC* is not responsive to vernalization (Finnegan et al., 2004).

The identification of putative *FLX* gene in gladiolus raises the question whether gladiolus follows an *Arabidopsis* model of the flowering pathway. In the winter annual *Arabidopsis thaliana*, flowering is promoted after vernalization, which suppress the floral suppressor *FLC* which is upregulated by *FRI*, through activation of *FRI* complex of (*FRI*, *FRL1*, *FRL2*, *FES1* and *SUF4*) proteins in addition to *FLX* protein. *FLX* was proven to provide transcriptional activity for the *FRI* complex (Choi et al., 2011). A loss of function of *FLX* in *Arabidopsis thaliana* resulted in early flowering phenotypes (Dennis et al., 2008), which indicates the clear role of *FLX* in flowering. The role of *FLX* in gladiolus has not been tested, particularly in RGC-1 genotypes and pedigrees; thus, *FLX* upregulation of *FRI* in gladiolus would be a rational approach. However, *FRI* was not detected in gladiolus, using the primer design of *Arabidopsis thaliana FRI* (At4g00650) because *FRI* gene was not detected in any monocotyledon species, thus the primer used is *Arabidopsis thaliana*, the test did not detect *FRI* gene in all 17 gladiolus genotypes without finding a single match (Aljaser, J. unpublished data). In addition, *VRN2*, the repressor of flowering in cereals and *A. thaliana* was not detected in gladiolus, using the primer design of *Triticum monococum*, *T. durum* and *Hordeum vulgare* (Aljaser, 2020, Appendices A1, A2 and A3). This is not conclusive evidence as the genetic similarities between *A. thaliana* and *Gladiolus* are low, given that *GhUFC-A* is ~32% and *GhFLX* is 50% identical to *A. thaliana* genes. Therefore, there could be an *FRI* gene in gladiolus but would require better primer design to locate the gene, because the presence of the *GhFLX* gene might indicate in the presence of other flowering repressor genes as *FRI* protein upregulates *FLX* in *Arabidopsis thaliana* and this is part of the flowering pathway (Choi et al., 2011). In addition, *Musa acuminata, Elaeis guineensis* and *Ananas comosus* are all monocots and tropical species which have *FLX* and *SUF4* genes which are part of *FRI* complex (Amasino and Michaels, 2010). This indicates the presence of some of *FRI* complex components while a lack of identification of *FRI* gene itself creates divergent possibilities: either there are *FRI* and *FLC* genes in these species or a lack of these genes and the presence of *SUF4* and *FLX* genes have other unknown flowering pathway purposes. Since *GhFLX* and *GdFLX* have similarities to *FLL1*, reaching up to 50% identity in amino acids, *FLX* gene is part of gene family, *FLX-LIKE1* (*FLL1*), *FLX-LIKE2* (*FLL*)), *FLX-LIKE3* (*FLL3*), *FLX-LIKE4* (*FLL4*) (Choi et al., 2011; Lee et al., 2013). While the role of *FLL1* in flowering pathway is not proven, *FLX* and *FLL4* are most crucial genes in control of flowering time in *Arabidopsis* (Lee and Amasino, 2013).

The relation between *UFC* and *FLX* is addressed in Dennis et al 2008, pointing that a mutation in *FLX* would influence in *UFC* expression, in which *UFC* expression is reduced in *flx* mutant in *A. thaliana*, this was a clear study to establish the influence of between the two genes.

The next step in this research would be to identify the *UFC* gene in diploid gladiolus species to determine if the allele is similar to *GhUFC-A, GhUFC-B* or a third different allele, using diploid species would simplify the study to determine the function of *UFC* protein in gladiolus by silencing and knocking out the gene. Locating the physical location of the *UFC* gene in *Gladiolus* will help in testing if there are other *UFC* genes in gladiolus as part of a *UFC* gene family, since the first discovered *UFC* gene (At5g10150) in *Arabidopsis thaliana* is located in the cluster genes *UFC, FLC* and *DFC* on chromosome 5 (Finnegan et al., 2004). *UFC* is also designated *SOK2* and the other *UFC* genes are grouped in *SOK* gene family such as *SOK1* (At1g05577), *SOK3* (At2g28150), *SOK4* (At3g46110) and *SOK5* (At5g59790) (Yoshida and Weijers et al., 2019).

In terms of the *FLX* gene, identifying the *FLX* gene in Eurasian species of *Gladiolus*, particularly *G. italicus*, *G. imbricatus* and *G. communis*, would be informative since these perennial species live in temperate habitats that require vernalization to break corm dormancy in winter season (Cohen and Barzilay, 1991; Cohat, 1993; Gonzalez, 1998). Conversely, identifying *FLX* in subtropical gladiolus species, such as *G. crassifolius*, *G. laxiflorus* and *G*. *atropurpureus* (Goldblatt, 1996), would allow the comparison of *FLX* among these different habitats may support the influence in *FLX* in the process of flowering. Furthermore, the use of transgene silencing of *FLX* in gladiolus would determine whether or not *FLX* influences the production of a rapid flowering phenotype gladiolus (RGC-1), as was reported in loss of function of *flx* in *A. thaliana* (Dennis et al., 2008). In conclusion, the discovery of *UFC* and *FLX* genes in gladiolus provides insight of the better understanding for flowering and vernalization response in ornamental geophytes.

## Conclusion

Rapid generation cycling is a powerful tool can be implemented to reduce the juvenility period in perennial crop and should be applied in breeding program ideotype. Although it is possible to annualize a perennial crop through genetically modifying the flowering pathway by overexpression a positive flowering regulator or inserting blocker of flowering suppressor (Srinivasan et al., 2012), yet those methods are biotechnological methods and require much regulation and approval, or sometimes rejected. Therefore, conventional breeding methods for early flowering are wildly accepted and does not require much regulation for releasing the cultivar.

Geophytes such as gladiolus are perennial and take 3 – 5 years to flower from seed; likewise, tulip and daffodil each take 4 – 6 years (Fortanier, 1973). The University of Minnesota gladiolus breeding program aims to reduce the number of years to reach flowering (Anderson et al., 2015) and successfully “annualized” the perennial gladiolus to flower from seed in the first year. Gladiolus breeding for rapid generation cycling was first accomplished by Roemer (Roemer, 1907) and was commercialized in market (Burpee^®^, 1913), Minnesota Gladiolus Breeding Program successfully breed for rapid generation cycling using different genotypes than Roemer. The RGC lines was superior than commercial cultivars, by reaching to flowering from seed in less than a year, petite height suitable for potted production without the need of using plant growth retardants to reduce the height for seed propagated gladiolus.

The use of plant growth retardants in floriculture crops is demanding to create uniformity, growth retardants are used regularly on potted production (Miller et al., 2012), the taller the crops is the higher the concentration used. Gladiolus is relatively tall flowering crops ranging 1-2 meters in height, reducing the height using plant growth retardants require higher concentration. However, high concentrations show inhibition to flowering as gibberellins is one of the factors of flowering (Ehrich, 2013; Kamenetsky, et al., 2012). corms of RGC genotypes able to reach flowering under the use of high GA-inhibitor concentration. In addition, RGC genotypes able to reach flowering under such inhibiting factors while maintaining a shorter stature proves the superiority of RGC lines to be candidate for potted production gladiolus. Another trait in RGC genotypes is their ability to re-sprout by having reduce dormancy, which break the typical requirement of vernalization in gladiolus.

Vernalization is the exposure of cold weather for time period then subsequent induction of flowering. Temperate crops require vernalization to flower (Dubcovsky, 2009). The lack of vernalization requirement in RGC genotypes can acts as a mutant in the studying of vernalization genes in gladiolus. These features add value of RGC genotypes from economic value and research value as well. In ornamental plants, flowering is a crucial step for the success of cut flower production. Therefore, understanding the flowering pathway and the gene expression is important for efficient selective breeding for rapid generation cycling. Furthermore, the identification of flowering genes in geophytes is poorly understood. An important gene in flowering is *FLOWERING LOCUS C* (*FLC*) which is a major flowering repressor found in *Arabidopsis* and many dicot species, *FLC* plays vital role in the control of flower initiation (Michaels and Amasino, 1999). However, the lack of identifying *FLC* in many monocot species and discovery of another mechanism of flowering in monocots, such as plant age in both maize and rice (Leeggangers et al., 2013), led researchers to suggest that monocot geophytes could also be *FLC-* independent for flower initiation. The search for *FLC* gene and its regulatory genes in gladiolus is a step to uncover the flowering pathway in geophytes. To uncover if *FLC* is present in Gladiolus, we searched for linked genes with *FLC*. In *Arabidopsis*, *FLC* is adjacent to two genes, *UPSTREAM OF FLOWERING LOCUS C* (*UFC*) and *DOWNSTREAM OF FLOWERING LOCUS C* (*DFC*). The both *UFC* and *FLC* genes are downregulated by vernalization *FLC* (Dennis *et al*., 2004). The discovery of *UFC* in gladiolus as well *FLOWERING LOCUS C EXPRESSOR* (*FLX*) is crucial to establish the flowering pathway, *UFC* gene being adjacent to *FLC* in *Arabidopsis* and responses to vernalization and *FLX* a gene that is upregulates *FRIGIDA* (*FRI*) which upregulates *FLC* is an early indicator in the presence of *FLC* homologue in gladiolus. These finds are important to understand flowering mechanism and gene of flowering pathway to utilize the knowledge in order to produce potted gladiolus either by using seed as germplasm or corm of RGC genotypes (Appendix; A4).

## List of Symbols and Abbreviations

DFC: DOWNSTREAM OF FLOWERING LOCUS C
FLC: FLOWERING LOCUS C
FLX: FLOWERING LOCUS C EXPRESSOR
FRI: FRIGIDA
UFC: UPSTREAM OF FLOWERING LOCUS C

## Acknowledgements

The authors wish to thank undergraduate students Allison Graper, Kaylie Niedzwiecki, Sarah Gardner, and Cora Rost and Research Scientists Michele Schermann and Rajmund Eperjesi for their help in growing the plants, helping with leaf sampling and DNA extractions.

## Data availability

*UFC* and *FLX* allelic gene sequences for the 17 gladiolus genotypes are deposited into NCBI (accession numbers forthcoming).

